# Early Classification of Motor Tasks Using Dynamic Functional Connectivity Graphs from EEG

**DOI:** 10.1101/2020.08.12.244921

**Authors:** Foroogh Shamsi, Ali Haddad, Laleh Najafizadeh

## Abstract

**Objective:** Classification of electroencephalography (EEG) signals with high accuracy using short recording intervals has been a challenging problem in developing brain computer interfaces (BCIs). This paper presents a novel feature extraction method for EEG recordings to tackle this problem.

**Approach:** The proposed approach is based on the concept that the brain functions in a dynamic manner, and utilizes dynamic functional connectivity graphs. The EEG data is first segmented into intervals during which functional networks sustain their connectivity. Functional connectivity networks for each identified segment are then localized, and graphs are constructed, which will be used as features. To take advantage of the dynamic nature of the generated graphs, a Long Short Term Memory (LSTM) classifier is employed for classification.

**Main results:** Features extracted from various durations of post-stimulus EEG data associated with motor execution and imagery tasks are used to test the performance of the classifier. Results show an average accuracy of 85.32% about only 500 ms after stimulus presentation.

**Significance:** Our results demonstrate, for the first time, that using the proposed feature extraction method, it is possible to classify motor tasks from EEG recordings using a short interval of the data in the order of hundreds of milliseconds (e.g. 500 ms).This duration is considerably shorter than what has been reported before. These results will have significant implications for improving the effectiveness and the speed of BCIs, particularly for those used in assistive technologies.

## 1. Introduction

Brain computer interfaces (BCIs) are designed to establish a communication link between the human brain and external devices. BCIs offer the possibility of generating commands from brain recordings to control external devices such as those used in assistive technologies. Among different types of BCIs, electroencephalography (EEG)-based BCIs have received considerable attention in the BCI research community due to their advantages including non-invasiveness, high temporal resolution, low-cost, and portability [1, 2].

Two important metrics that impact the performance of BCIs and influence their efficiency and practicality in various applications are 1) the buffering lag, *i*.*e*. the duration of recordings required by the classification algorithm to create commands (which largely determines the speed of the BCI), and 2) the classification accuracy (which determines the reliability of the BCI). Clearly, an algorithm capable of classifying a number of different tasks with high accuracy using a “short” interval of recordings is of great interest.

In EEG-based BCIs, motor imagery (MI) tasks, targeting imagination of different forms of muscle movements, such as arms, hands, tongue, and legs have been commonly used. To this date, numerous classification algorithms, utilizing various feature extraction and classification techniques have been used in EEG-based BCIs for decoding motor execution/imagery tasks [3]. To provide an overview of recent work in this domain, in Table 1‡, a comparison of recent work is given. These studies have been categorized into different groups based on the buffering lag, which is the duration of the data that needs to be collected before a prediction can take place. We divided these studies into three groups of 1 − 2 s, 2 − 4 s, and equal or greater than 4 s. For each study, we have summarized information on the number of classes, number of EEG electrodes (channels), feature extraction domain (time, frequency, time-frequency), classification algorithm, and reported classification accuracy.

**Table 1:**
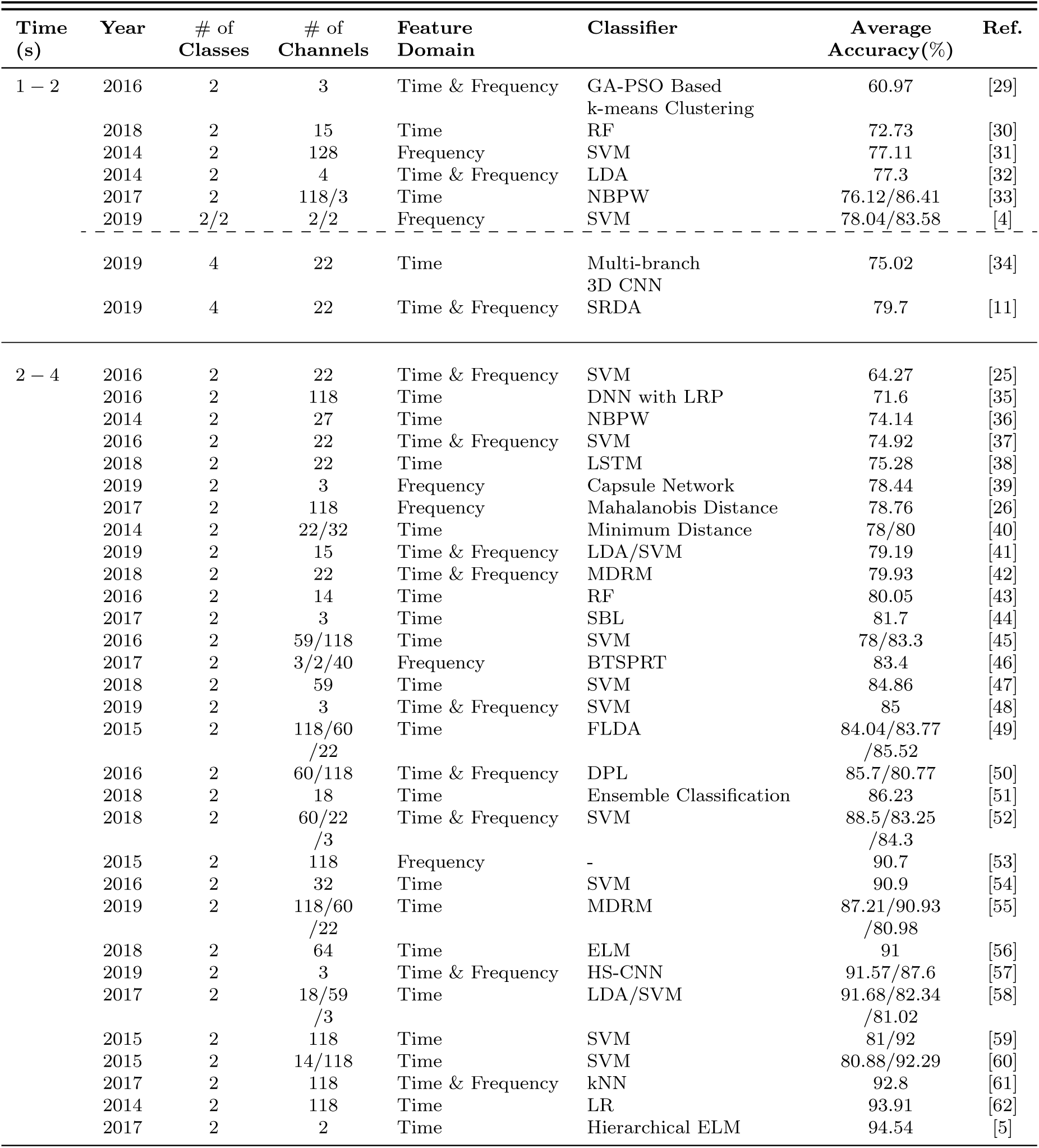
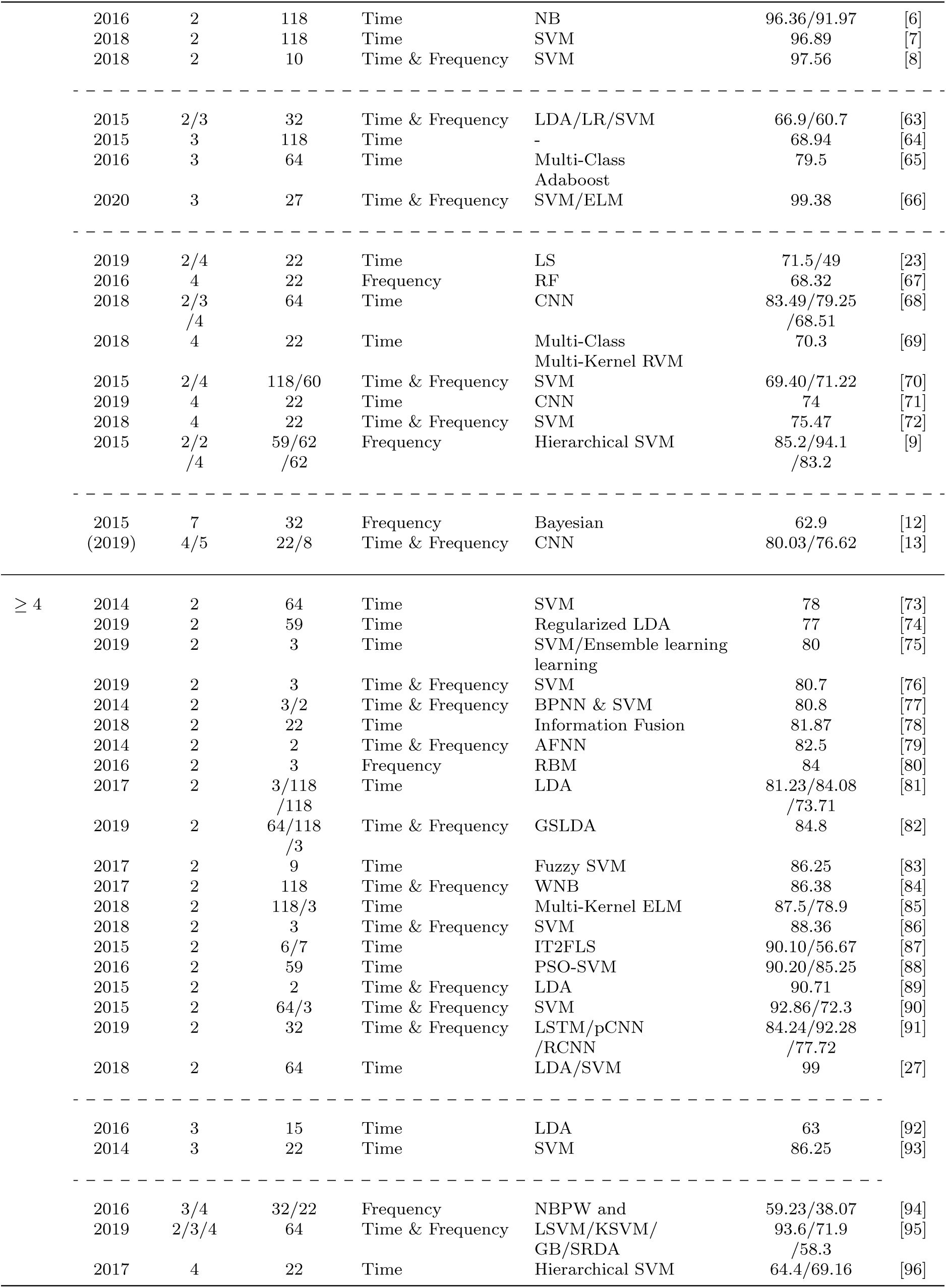
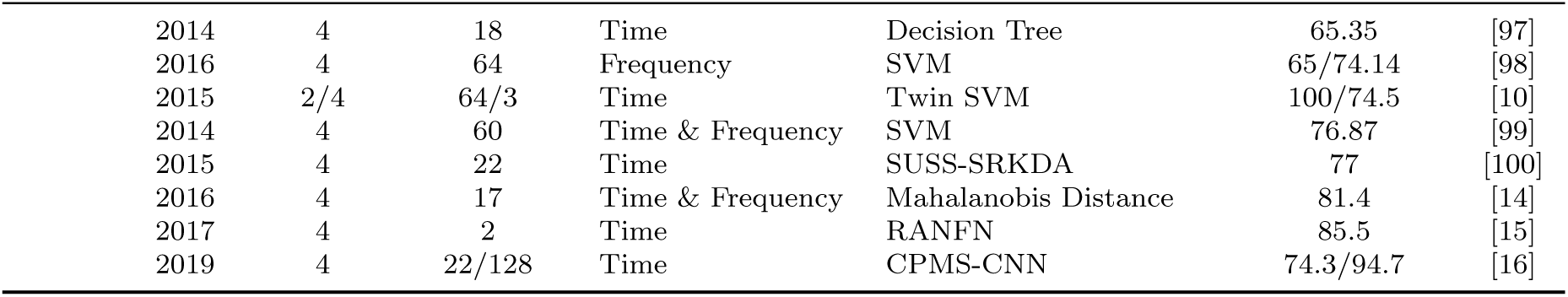
Performance comparison of recent studies for classification of MI tasks in EEG-based BCIs.

From Table 1, one can see that the majority of studies have considered two-class problems. For two class problems, the highest accuracy results for using 1 − 2 s of data were reported in [4], for using 2 − 4 s of data in [5, 6, 7, 8, 9], and for using equal or greater than 4 s of data in [10]. For multi-class problems, the highest accuracy results were achieved in [11] for using 1 − 2 s of data, in [9, 12, 13] for using 2 − 4 s of data, and in [14, 15, 16] for using equal or greater than 4 s of data. Considering these studies, it can be concluded that with respect to the required buffering lag, the decoding process has been relatively slow. This slowness causes challenges in BCI applications, where a sequence of commands needs to be decoded and executed in order to complete a task, such as moving a robotic arm from one place to another. There is therefore, a crucial need to develop new methods capable of early decoding of EEG signals to improve the efficiency of BCIs. In this paper, we aim to target this problem, and propose a framework, which requires a relatively short duration of EEG data (in the order of hundreds of milliseconds) to achieve high accuracy in classification.

Our proposed approach is based on functional connectivity and the hypothesis that for the execution of tasks, the brain relies on dynamic interactions among its different regions [17, 18, 19, 20]. Previous work [21, 22, 23, 24] have indicated that brain’s functional connectivity throughout the course of movement-related (execution/imagery) tasks can be used to study the underlying mechanisms involved in performing those tasks. These results suggest the potential of acquiring useful information from functional connectivity networks for discriminating various motor-related tasks. Indeed, functional connectivity has been utilized in recent EEG-based BCI work. As an example, in [25], phase-locking value (PLV) was used to estimate functional connectivity patterns from EEG recordings. The authors then identified node-pairs in constructed functional networks that discriminate resting state vs motor imagery tasks. Using this information along with a subject-specific frequency band selection procedure, features were extracted to discriminate left and right hand motor imagery tasks, where an average classification accuracy of 64.27% was achieved. In another connectivity-BCI study [26], after identifying functional connectivity patterns using partial directed coherence metric, a statistical selection method based on the appearance rate of directed connectivities and larger partial directed coherence magnitude was used to characterize task-specific functional connectivity networks. The identified networks were then used as features and an average classification accuracy of 78.76% for discriminating right hand vs right foot motor imagery tasks was reported. In [27], the feasibility of using functional connectivity patterns for classification of motor imagery (left vs right hand) was explored. In this study, interactions among EEG electrodes were modeled as graphs which were constructed using motif synchronization method, and then, various graph metrics were used as features for the classification problem. The results of this study demonstrate the feasibility of using graph methods for the classification of motor imagery tasks. In another work [23] graph measures were employed as features for decoding motor imagery tasks, resulting in an average classification accuracy of 71.5% for classifying pairs of left hand, right hand, foot, and tongue motor imagery tasks. In [11], connectivity graph measures combined with channel-based time/frequency domain features were employed for the classifications of motor imagery tasks, and an average accuracy of 79.69% was reported for a 4-class motor imagery classification. These studies demonstrate the feasibility of using functional connectivity networks for decoding motor-movement tasks in BCIs. However, as seen in Table 1, in these functional connectivity-based BCI studies, features were extracted from durations of equal or greater than 2 s. As for dynamics, dynamic brain functional networks corresponding to motor imagery tasks during the event-related synchronization (ERS) and de-synchronization (ERD) periods were investigated in [22], where it was shown that different stages of motor preparation and imagery can be characterized by network measures. Their findings demonstrated that the underlying dynamic information processing during motor imagery tasks can be described by dynamic functional connectivity networks, suggesting the potential for using dynamic functional connectivity in BCIs. In [28], empirical mode decomposition phase locking method was used to model the functional connectivity between EEG channels, and time-frequency connectivity maps were generated by computing time-dependent mean clustering coefficients of graph nodes for different frequency bands. A Hidden Markov Model (HMM) was then employed to classify the dynamics of the mean clustering coefficients in different frequency bands, indicating that the dynamics of functional connectivity networks could provide useful information for differentiating motor tasks.

Motivated by the results of functional connectivity-based BCIs, in this paper, we propose a new framework that utilizes the spatial and temporal (dynamic) characteristics of brain functional networks for early decoding of motor execution and imagery tasks. Our hypothesis is that for different movement execution/imagery tasks, the “*spatial* “distribution of the brain functional networks as well as their “*dynamics*” provide discriminatory information for the classification problem, even within a short interval after the task onset.

In the proposed framework, EEG recordings are first segmented using our recently-proposed segmentation method [101, 102, 103] to identify the time intervals (segments) during which the spatial distribution of the underlying functional networks stays quasi-stationary. Functional connectivity networks for each identified segment are then localized to generate functional connectivity graphs. The resulting graphs, extracted from the sequences of the segments, are then vectorized and passed to the classifier. Considering the dynamic nature of the extracted functional connectivity graphs, we employ a long short-term memory (LSTM) network as the classifier, which allows for using the information in sequence. To evaluate the early decoding capability of the proposed framework, we use the sequence of functional connectivity graphs extracted from the corresponding sequence of identified post-stimulus EEG segments, starting with the very first segment. We also explore changes in the decoding accuracy as a result of increasing the duration of EEG recordings used for classification. It is worth mentioning that the proposed method is a synchronous BCI algorithm.

Two datasets are used to evaluate the performance of the proposed framework. The first dataset is collected in our lab and includes various forms of tongue/hand motor execution and imagery tasks [104]. The reason for choosing tongue movement execution task is that voluntary tongue movement is mostly preserved in patients with severe motor disorders, such as those with high spinal cord injury [105, 106], and therefore, can be utilized in BCIs to increase the number of commands in BCI-based assistive technologies for these patients. The second dataset is taken from a publicly-available source (BCI competition IV, dataset IIa) and comprises four different imagery tasks [107]. Our results demonstrate that although classification accuracy generally improves over time, an average accuracy of 85.32% can be reached only 500 ms after task onset. In summary, the contributions of this paper are as follows: 1) we present a new framework that utilizes the spatial and dynamic characteristics of brain functional networks for decoding motor tasks from EEG recordings; 2) we demonstrate, for the first time, that it is possible to achieve reasonable classification accuracy using only a short interval of the data in the order of hundreds of milliseconds (e.g. 500 ms) after the task onset, which can pave the path to further enhance the performance of the BCIs in terms of the required buffering lag and increasing the decoding speed; and 3) we consider classification of tongue movement execution versus hand/tongue movement imagery, which offers the possibility of increasing the number of commands in BCIs.

The rest of the paper is organized as follows: the proposed methods for feature extraction and classification of EEG data are presented in Section 2. In Section 3, the experimental paradigms and data collection procedures are described. Classification results and discussions are presented in Sections 4 and 5, respectively, and the paper is concluded in Section 6.

## 2. Proposed Method

A functional network corresponds to a set of brain regions that exhibit correlation to a common time-course, suggesting that they are collaborating functionally when the brain is at rest or when it is engaged in executing a task [108]. It is also now known that functional networks are dynamic, *i*.*e*., they sustain their inner-connectivity for short intervals of time [17, 18]. As will be discussed here, our proposed novel feature extraction/classification framework uses functional connectivity networks as well as their dynamics, to enable the possibility of early decoding of EEG signals. An overview of the proposed framework is illustrated in Figure 1. We consider a LSTM classifier to take advantage of the information contained in the dynamics of the extracted features. In what follows, we provide details for each step in the proposed framework.

**Figure 1:**
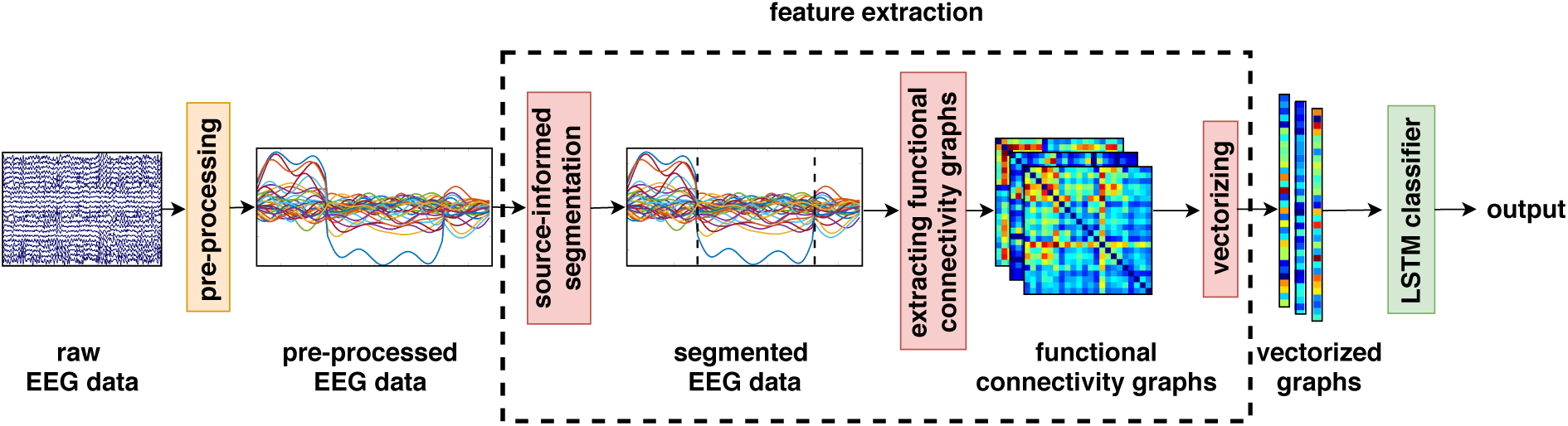
Overview of the proposed framework.

### 2.1. Proposed Feature Extraction Framework

The proposed feature extraction framework is comprised of two main steps: first, the pre-processed EEG data is segmented into quasi-stationary intervals, and then functional networks in each segment are localized and the corresponding graphs are constructed, which will then be used as features for the LSTM classifier.

#### 2.2.1. EEG Segmentation

The first step of feature extraction uses our recently developed source-informed segmentation algorithm [101]. The main objective of this step is to find intervals in the EEG recordings during which “*the spatial distribution*” of the cortical functional networks stays quasi-stationary. In other words, this step detects the time points in the EEG data, at which there are changes in the location of cortical networks. Therefore, this segmentation approach provides a functionally-relevant means for segmenting EEG data, which is different from the commonly-practiced model-based (e.g. autoregression) and metric-based (e.g. change point detection) segmentation approaches.

The details of the source-informed segmentation algorithm is presented in [101], and the algorithm was utilized in [109, 102, 103, 110, 111]. Briefly, this segmentation technique employs singular value decomposition (SVD) along with a reference/sliding window approach to identify targeted time intervals in the EEG data. In [101] we have proved that the most significant left singular subspace of the EEG data captures the “*spatial locality* “features of the cortical functional networks and can be used as a feature space in the segmentation algorithm. Therefore, a significant change in this span of the feature space, along the time axis, would indicate a change in the spatial distribution of the cortical functional networks. Using a reference/sliding window approach, this feature space can be dynamically extracted. The segment boundaries are detected by statistically comparing the residual error resulting from projecting the block of EEG data matrix under a reference window, on one hand, and that under a sliding window, on the other hand, onto the feature subspace [101]. It is worth mentioning that this segmentation algorithm is performed without using source localization methods, and uses the information from all EEG channels to detect the boundaries of the segments.

#### 2.1.2. Functional Connectivity Graphs

The second step of the feature extraction involves identifying the functional networks sustaining their connectivity during each identified segment and calculating the corresponding graphs. Since the nodes of these networks are taken to be the EEG electrodes, the problem of volume conduction needs to be addressed [112, 113]. For this purpose, the surface Laplacian operator [114, 115] is applied to the EEG data in order to minimize channel coupling. The segment-wise average value of each EEG channel is then removed, in order to allow the next step to focus on the temporal patterns of the activities, rather than the intensities, of these channels.

Towards identifying the functional networks forming during a given segment, we first note that each of these networks is characterized by a distinct time-course with which the involved nodes show high correlation. Here, we use low-rank SVD to recover a minimum mean square error (MMSE) estimate of the zeros-mean channel activities conforming to these time-courses, thus, to their associated networks. For more details about this approach refer to [102]. The functional connectivity graph that captures the networks forming during that segment can finally be calculated as the correlation matrix among the estimated channel activities.

To illustrate an example of the outcome of the proposed methods, we applied the proposed segmentation and functional connectivity graph extraction algorithms to 4 randomly selected trials from the BCI Competition IV-IIa dataset (see Section 3 for more details about the experimental paradigm). Each trial was taken from one class of motor imagery (left hand, right hand, both feet, and tongue). The results for 110 ms are shown in Figure 2 as examples. It can be seen that, the length and the number of the segments, as well as the extracted graphs are different for each class, further suggesting that these features can be utilized to differentiate various forms of motor-movement classes.

**Figure 2:**
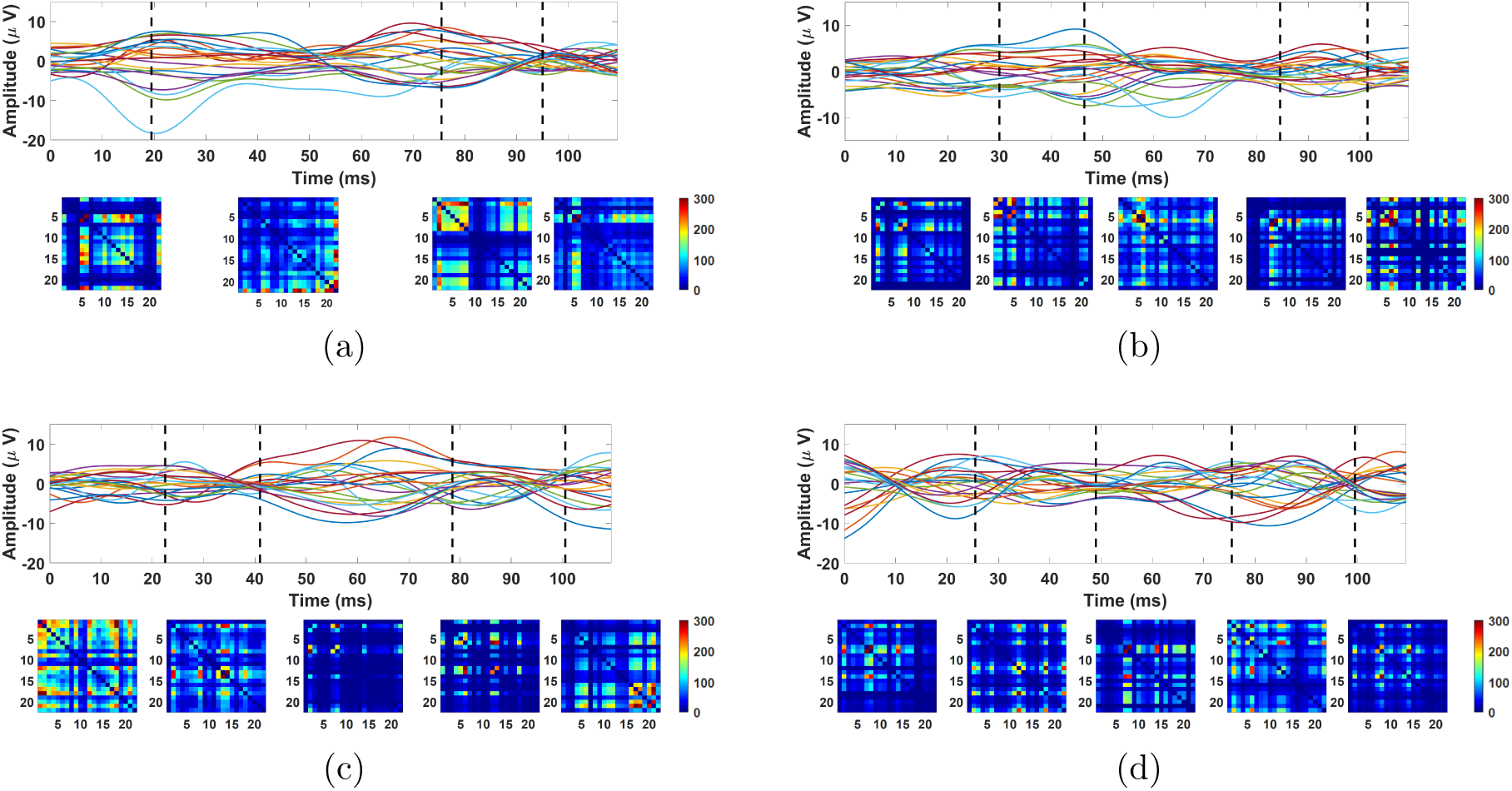
Segmentation and extracted color-coded functional connectivity matrices for each segment of randomly selected trials from (a):left hand motor imagery, (b): right hand motor imagery, (c): both feet motor imagery, and (d): tongue motor imagery of BCI Competition IV-IIa dataset. As can be seen, across classes of motor movement tasks, there are variations in the number and the duration of identified segments, as well as in patterns of constructed graphs.

### 2.2. Classification Scheme

Due to the dynamic nature of the brain, it is important to consider the information contained in the temporal sequence of the extracted features from identified segments. We address this point by employing a LSTM classifier. The input to these classifiers are variable-length temporal sequences of connectivity graph-based vectors. The variation in the lengths of these sequences comes from the variation in the number of the identified segments, spanning any fixed time interval.

The structure of the classifier is shown in Figure 3. A neural network with three hidden layers is used: a fully-connected layer consisting of 20 neurons, an LSTM layer consisting of 20 neurons with a single-step delay feedback loop around the second hidden layer, and another fully-connected layer consisting of 2 neurons. These layers are followed by *softmax* and classification layers. An overall classification decision is made by the neural network after it processes each segment in the sequence of dynamic functional connectivity graphs.

**Figure 3:**
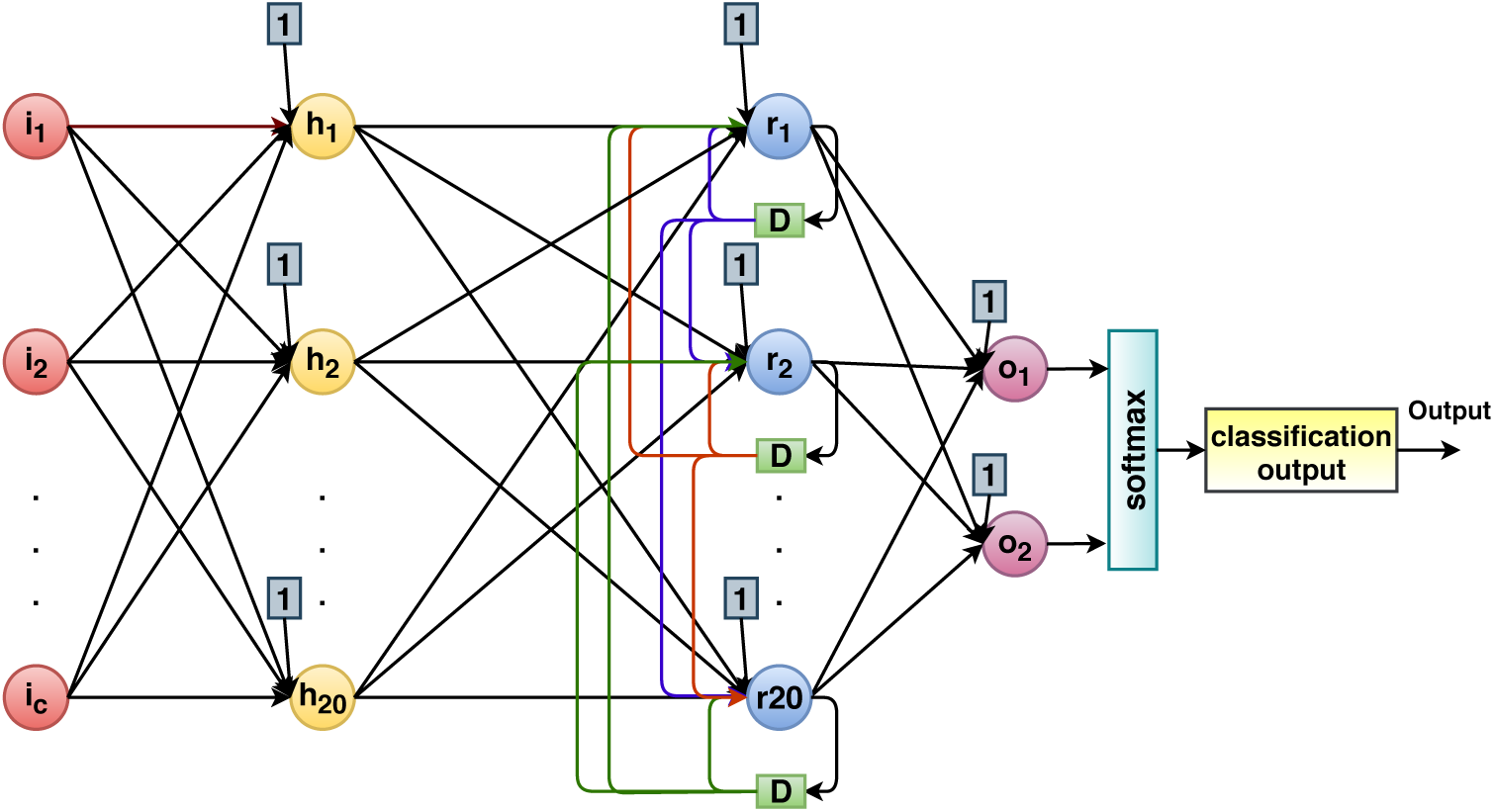
The structure of the LSTM classifier.

## 3. Experimental Procedure

To evaluate the performance of the proposed method, two datasets have been used. The first dataset was collected in our lab and the second dataset was available from the BCI Competition IV-IIa. Detailed descriptions of the experimental paradigms for each dataset are provided in this section.

### Dataset 1

Eight healthy, right-handed volunteers (5 males and 3 females) aged between 20 and 35, participated in the study. Written informed consents approved by Rutgers’ Institutional Review Board (IRB) were obtained prior to experiments. Subjects were seated in a comfortable chair with a display in front of them as shown in Figure 4. The experiment included 3 blocks of dictated motor execution tasks and 3 blocks of dictated motor imagery tasks separated by short breaks. On average each block took approximately 7.5 minutes. In execution/imagery blocks, subjects were instructed to *move/visualize moving* their tongue upward or downward or *squeeze/visualize squeezing* their left or right hand if they saw an arrow pointing up, down, left, or right, respectively. The direction of movement, using an arrow, was shown for 1 s, followed by a diamond stimulus which was shown for 2 s. Subjects were instructed to perform motor execution or imagery only after they saw the diamond stimulus and to continue until it disappeared. The inter-trial interval was set to 2 − 4 s (see Figure 5(a)). In each block, 15 trials of each class were performed (a total of 45 trials per class). EEG data was collected using a 128-channel EEG system (Brain Products), at a rate of 2000 samples/sec. We used 32 EEG channels for data analysis. The selected channels are shown in Figure 5(b). The electrodes were positioned based on the international extended 10 − 20 electrode placement system.

**Figure 4:**
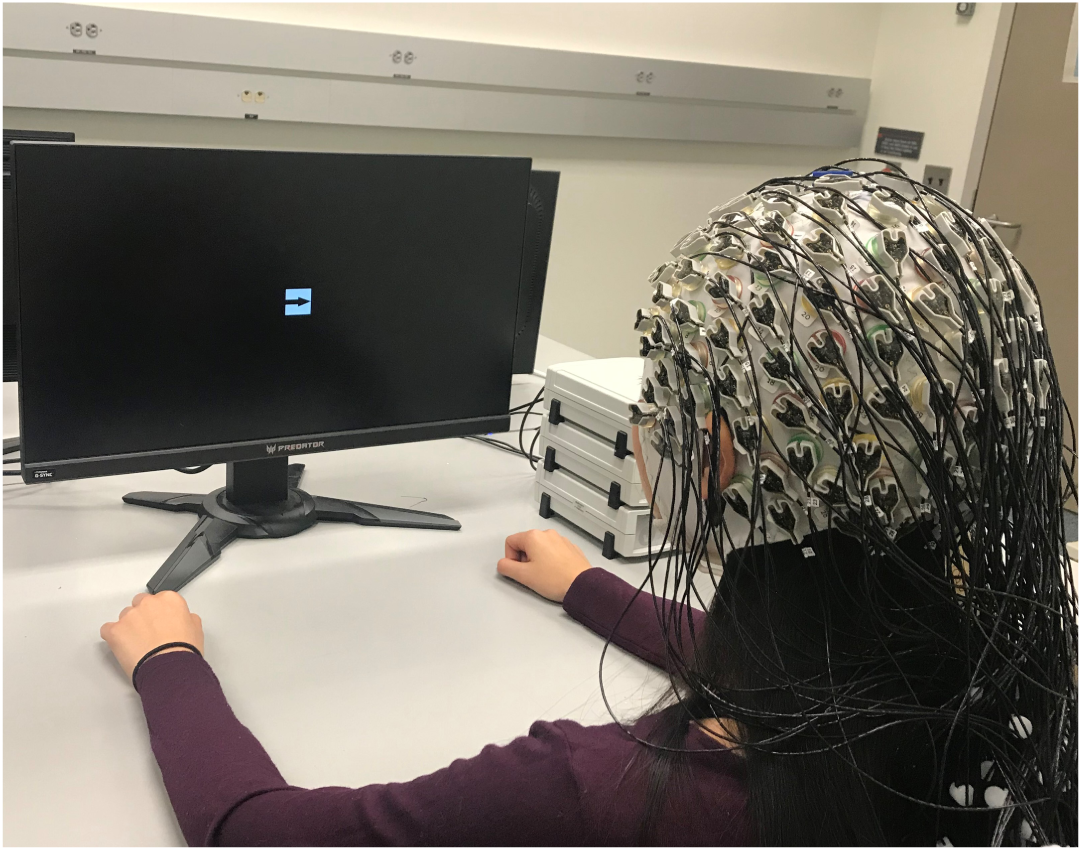
Experimental setup for Dataset 1.

**Figure 5:**
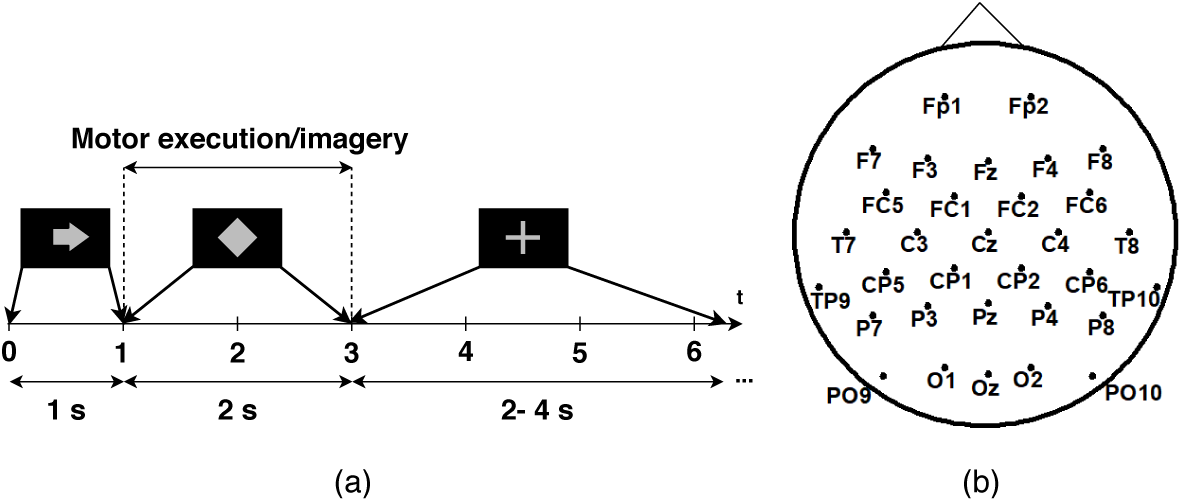
Dataset 1-(a): Timing scheme of a single trial. (b): EEG electrodes montage.

The 2 s EEG data obtained during the presentation of the diamond stimulus, was extracted from each trial and was preprocessed using EEGLAB toolbox [116]. The data was filtered using a band-pass finite impulse response (FIR) filter in the range of 1 to 50 Hz. All imagery and execution trials were treated the same for preprocessing. The artifacts were then removed using visual inspection of the trials and independent component analysis (ICA).

### Dataset 2

This online dataset was used to conduct a performance comparison with existing work (see Section 5). The dataset provided by the Laboratory of Brain-Computer Interfaces, Graz University of Technology have been used in several BCI papers. The EEG data was collected from 9 right-handed subjects using 22 Ag/AgCl electrodes (Figure 6(b)) and 3 EOG channels, at a rate of 250 Hz. The data was collected in two sessions on different days. Each session comprised 6 runs with short breaks between blocks. During each run, the subjects were asked to perform four motor imagery tasks including the imagination of movement of the left hand, right hand, both feet, and tongue. At the beginning of each trial, a fixation cross is shown on the black screen and a short acoustic warning tone is presented. After 2 sec, an arrow pointing to the left, right, down or up, corresponding to the motor imagery of left hand, right hand, both feet, and tongue, appears on the screen for 1.25 s, followed by a fixation cross. Subjects were asked to perform motor imagery task until the fixation cross disappears at *t* = 6 s. A short resting time was considered before starting a new trial. Each run consisted of 12 trials of each class, yielding a total of 72 trials per class in each session. A visual illustration of a single trial is shown in Figure 6(a).

**Figure 6:**
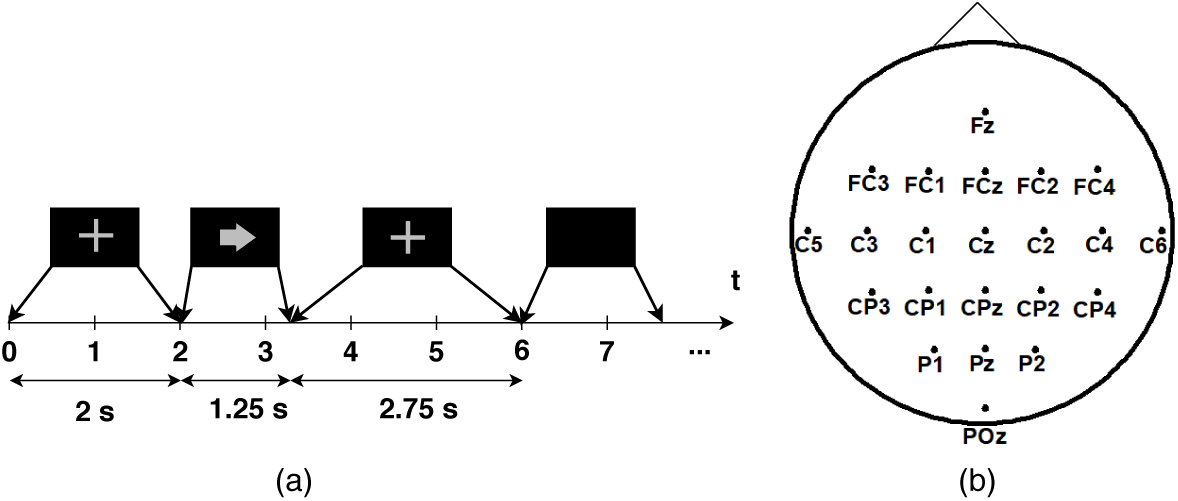
Dataset 2-(a): Timing scheme of a single trial. (b): EEG electrodes montage.

Previous studies that used this dataset, have mostly considered the [3 − 6] s window as the interval during which motor imagery tasks are performed [37, 42, 23, 25, 52, 69, 72] for the analysis. Therefore, we also extracted this 3-s motor imagery interval from each trial. Data was then preprocessed in EEGLAB toolbox [116], using a band-pass FIR filter in the range of 1 to 45 Hz. For artifact removal, the trials marked as bad trials in the file that accompanied the data, were removed in the pre-processing step before removing artifacts via ICA.

Table 2 summarizes the total number of available and used trials for each dataset.

**Table 2:** Number of available and used trials for Dataset 1 and 2.

|  | Available trials | Used trials |
| --- | --- | --- |
| **Dataset 1** | 2160 (270 per subject, 45 per task) | 2160 |
| **Dataset 2** | 5184 (576 per subject, 144 per task) | 4699 |

## 4. Results

In this section, we present the results for the performance of the proposed feature extraction/classification framework for different durations of post-stimulus EEG data. The trials were separated into three randomized groups for: training (75%), validation (10%), and testing (15%) the classification model. For Dataset 1, features extracted from 2 s post-stimulus intervals were used for training and validation. However, for testing, different time intervals of the data were used to study the early decoding capability of the proposed framework.

Figure 7 displays the probability mass function (PMF) of the length of the identified segments for the duration of motor execution/imagery tasks across all subjects, tasks, and trials for Dataset 1. The average length of the segments is 22.2 ms with standard deviation of 8.7. The sequences of graph patterns from identified segments were passed to the classifiers and the decisions were made at the end of each segment. As the segments for different trials have different durations and ending points, to calculate the accuracy at any time instant, decisions were grouped based on the ending point of the corresponding segment into non-overlapping time bins of 100 ms duration.

**Figure 7:**
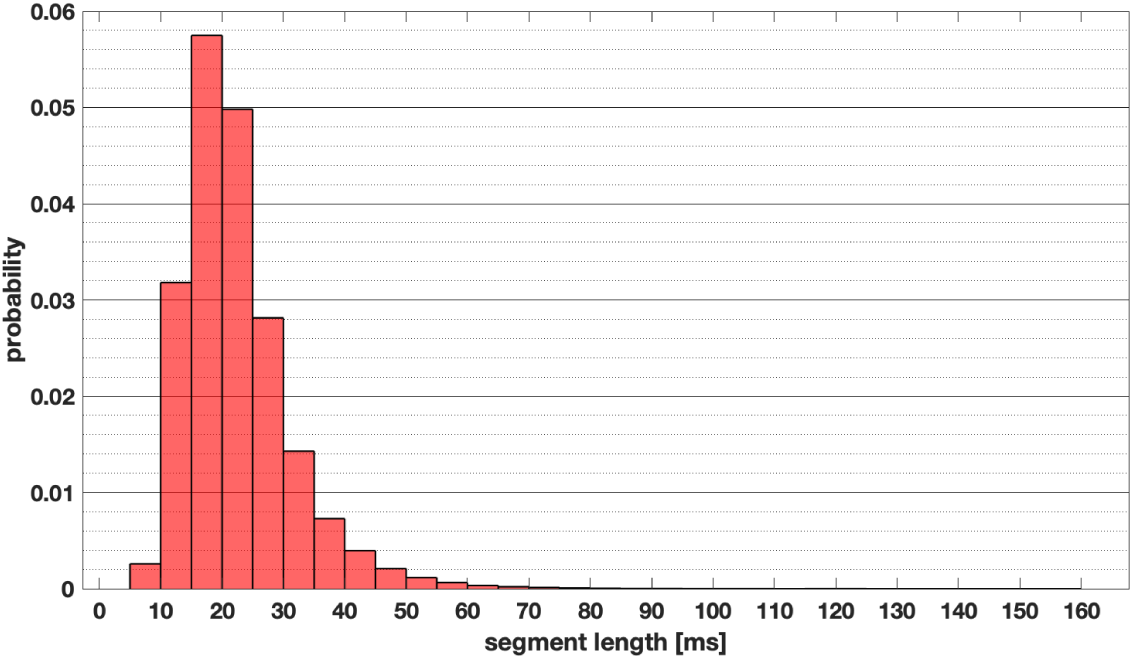
PMF of the segment lengths across all subjects, tasks, and trials. The average length of the segments is 22.2 ms with standard deviation of 8.7.

Let us describe this procedure with an example. For a randomly selected trial (e.g. subject 6, tongue imagery movement towards up direction, trial 9), the identified segments for the duration of [0 − 300] ms are as follows: [0 − 20.5] ms, [20.5 − 33.5] ms, [33.5 − 60] ms, [60 − 84] ms, [84 − 108] ms, …, [184 − 206.5] ms, [206.5 − 223] ms, [223 − 238] ms, [238 − 250.5] ms, [250.5 − 268.5] ms and [268.5 − 285] ms. The functional connectivity feature extracted from an individual segment is passed to the dynamic classifier to generate a single decision about the type of the task. The sequence of segments results in a sequence of decisions. Now, to calculate the average accuracy for example for the time bin [200 − 300] ms, the classification decisions from all segments whose end-point lies within the [200 − 300] ms bin are considered. In this example, these segments are [184 − 206.5] ms, [206.5 − 223] ms, [223 − 238] ms, [238 − 250.5] ms, [250.5 − 268.5] ms and [268.5 − 285] ms. Note that we did not include the decision from segment [285 − 304.5] ms in the calculation of accuracy here, because this segment does not end within the [200 − 300] ms bin.

### 4.1. Dataset 1 Classification Accuracy Using Short Duration of Post-stimulus EEG Data

For Dataset 1, 32 EEG channels (Figure 5) were used for the classification. For the segmentation step, 20 samples for both reference and decision windows were considered, which according to [101], it is a proper choice in terms of the performance of the algorithm in detecting segment boundaries. Using the 32 EEG channels, the size of the extracted functional graph matrices is 32 × 32. Since the extracted graphs are symmetric and include no self-loops, only the upper triangle from each graph matrix was extracted and put into a vector of size 496 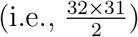. The extracted graphs for an exemplary subject and two randomly-selected trials are shown in Figure 8. One can see, qualitatively, that across tasks, there are differences in the patterns of the connectivity graphs as well as in the duration of the EEG data for which the graphs were constructed as determined by the segmentation algorithm.

**Figure 8:**
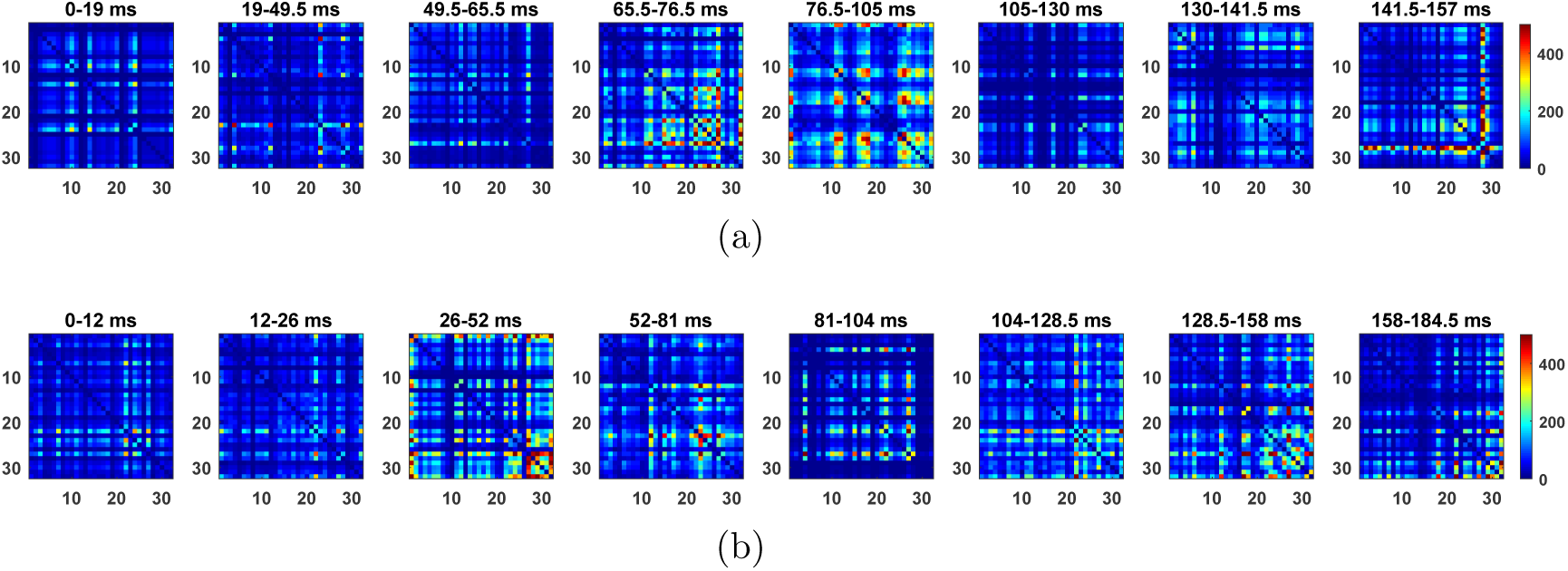
Extracted first 8 graph sequences of two randomly-selected trials from Dataset 1 for an exemplary subject (subject 6) (a): execution of tongue movement in the down direction, (b): imagery of right hand squeezing.

In this work, we focus on the classification of tongue movement execution vs tongue/hand imagery tasks. Selecting these pairs of motor execution vs imagery tasks was motivated by the fact that the tongue movement ability is often preserved in patients with motor disabilities, and its inclusion in BCIs can increase the number of control commands which is crucial in BCI-based assistive technologies. In contrast, patients in need of BCI technology usually do not possess hand movement execution ability. Therefore, discrimination of hand motor execution tasks vs tongue/hand motor imagery tasks was not considered in this study.

The accuracy, using the first 500 ms of the EEG data for classifying tongue movement execution vs tongue imagery (in various directions), and for tongue movement execution vs hand movement imagery (in various directions) for all subjects, and averaged across subjects are summarized in Tables 3 and 4, respectively.

**Table 3:** Classification accuracy results for tongue movement execution (TE) in up (U) or down (D) directions vs tongue imagery (TI) in up (U) or down (D) directions, 500 ms after the task onset.

|  | U (TE) vs U (TI) | U (TE) vs D (TI) | D (TE) vs U (TI) | D (TE) vs D (TI) | Average |
| --- | --- | --- | --- | --- | --- |
| **Subject 1** | 83.60 ± 9.59 | 79.93 ± 10.19 | 87.01 ± 8.25 | 80.90 ± 9.45 | 82.86 ± 9.39 |
| **Subject 2** | 84.86 ± 8.21 | 79.68 ± 10.27 | 78.95 ± 11.13 | 82.74 ± 9.69 | 81.56 ± 9.88 |
| **Subject 3** | 92.02 ± 6.68 | 94.60 ± 6.89 | 89.63 ± 8.24 | 90.12 ± 9.60 | 91.59 ± 7.94 |
| **Subject 4** | 93.61 ± 5.96 | 87.59 ± 7.38 | 94.62 ± 6.83 | 89.68 ± 8.19 | 91.37 ± 7.14 |
| **Subject 5** | 98.58 ± 5.67 | 98.70 ± 2.34 | 98.55 ± 3.14 | 98.02 ± 3.12 | **98.46 ± 3.78** |
| **Subject 6** | 91.52 ± 8.22 | 90.88 ± 6.99 | 81.37 ± 9.58 | 77.25 ± 10.27 | 85.26 ± 8.86 |
| **Subject 7** | 55.87 ± 11.24 | 50.61 ± 12.85 | 58.64 ± 11.35 | 53.31 ± 12.18 | 54.36 ± 11.92 |
| **Subject 8** | 77.45 ± 11.32 | 69.65 ± 12.71 | 77.09 ± 11.42 | 66.48 ± 12.98 | 72.67 ± 12.13 |
| **Average** | 84.69 ± 8.61 | 81.33 ± 9.30 | 83.23 ± 9.13 | 79.81 ± 9.84 | **82.27 ± 9.23** |

**Table 4:** Classification accuracy results for tongue movement execution (TE) in up (U) or down (D) directions vs imagery (HI) of left (L) or right (R) hands, 500 ms after the task onset.

|  | U (TE) vs L (HI) | U (TE) vs R (HI) | D (TE) vs L (HI) | D (TE) vs R (HI) | Average |
| --- | --- | --- | --- | --- | --- |
| **Subject 1** | 85.42 ± 10.62 | 85.80 ± 10.03 | 86.20 ± 7.85 | 84.13 ± 8.55 | 85.39 ± 9.33 |
| **Subject 2** | 75.48 ± 11.01 | 79.45 ± 10.42 | 69.95 ± 9.31 | 78.34 ± 10.48 | 75.81 ± 10.32 |
| **Subject 3** | 93.09 ± 7.68 | 94.94 ± 5.85 | 88.33 ± 8.93 | 91.19 ± 9.73 | 91.89 ± 8.18 |
| **Subject 4** | 95.12 ± 5.37 | 93.07 ± 6.93 | 94.05 ± 6.77 | 90.85 ± 9.17 | 93.27 ± 7.19 |
| **Subject 5** | 98.15 ± 3.31 | 98.07 ± 5.92 | 97.57 ± 4.38 | 99.14 ± 2.48 | **98.23 ± 4.22** |
| **Subject 6** | 93.98 ± 6.35 | 93.05 ± 7.47 | 85.30 ± 9.81 | 82.91 ± 9.38 | 88.81 ± 8.37 |
| **Subject 7** | 74.88 ± 10.60 | 85.68 ± 9.82 | 75.69 ± 11.31 | 80.37 ± 11.51 | 79.16 ± 10.83 |
| **Subject 8** | 74.301 ± 12.18 | 72.69 ± 11.26 | 67.65 ± 12.09 | 65.52 ± 12.31 | 70.04 ± 11.97 |
| **Average** | 86.30 ± 8.90 | 87.84 ± 8.70 | 83.09 ± 9.10 | 84.06 ± 9.62 | **85.32 ± 9.09** |

As expected, the accuracy results are subject-dependent. For Subject 5, the average accuracy for all classes was higher than 97%, while for subjects 1, 3, 4, and 6, the average accuracy results for different classes were above 80%. However, the average accuracy for tongue movement execution vs tongue imagery for subject 7 was around the chance level, which could be an outlier. Overall, after 500 ms of task onset, the average accuracy results of 82.27% and 85.32% were achieved for tongue movement execution vs tongue imagery, and tongue movement execution vs hand movement imagery tasks, respectively, across all subjects. These results show that the discrimination between tasks can be obtained within a short duration (in the order of hundreds of milliseconds) of task onset using the proposed framework.

### 4.2. Dataset 1: Effects of Increasing the Time Duration of the EEG Data on the Classification Accuracy

To this point, we only considered average classification accuracy 500 ms after task onset. To further study the effects of the post-task onset time elapse, on the accuracy results, the classification accuracy results for tongue movement execution vs tongue imagery, and for tongue movement execution vs hand movement imagery are plotted as functions of time in Figures 9 and 10, respectively.

**Figure 9:**
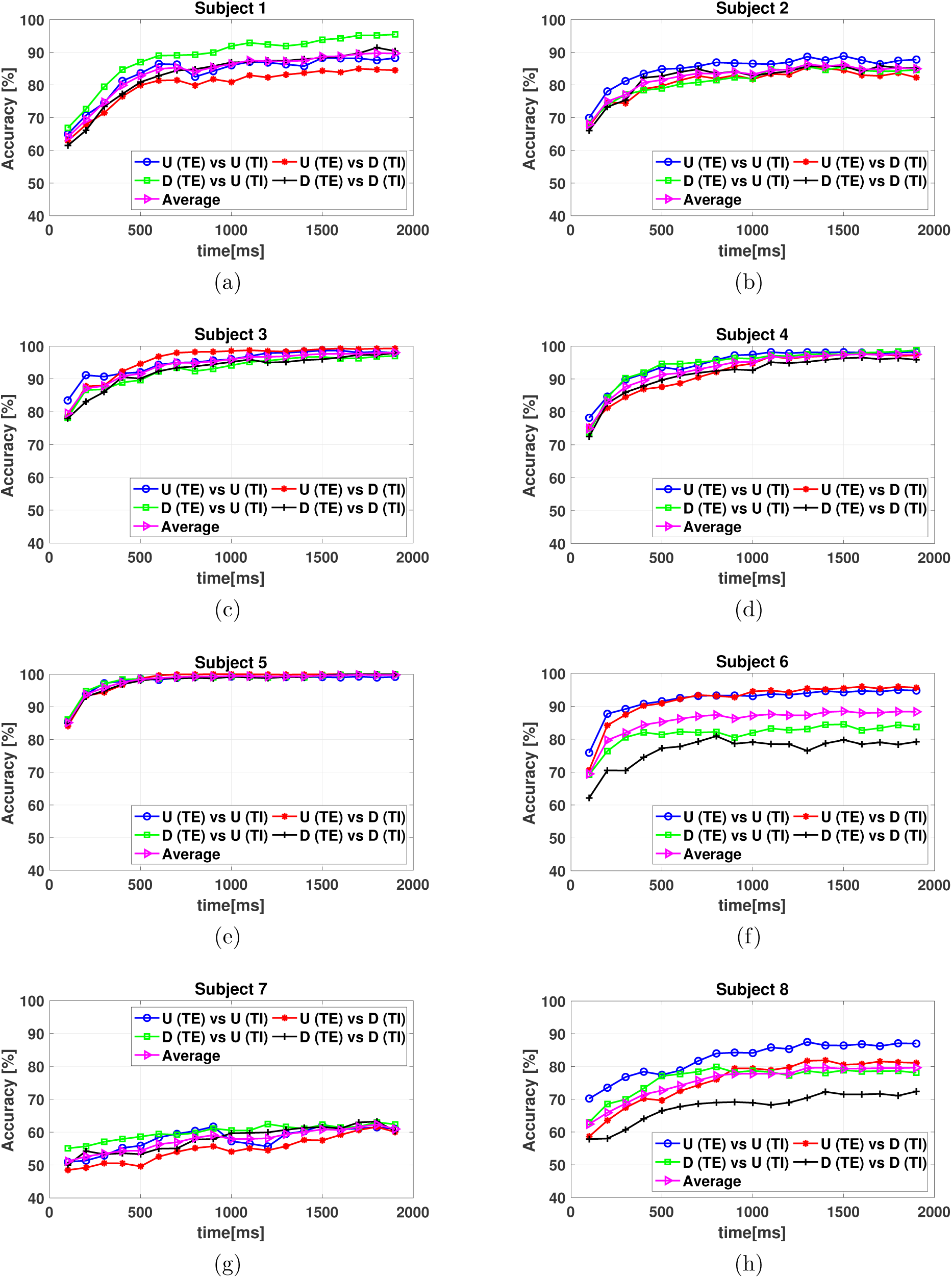
(a)-(h): Classification accuracy of tongue movement execution (TE) in up (U) or down (D) directions vs tongue imagery (TI) in up (U) or down (D) directions for all subjects.

**Figure 10:**
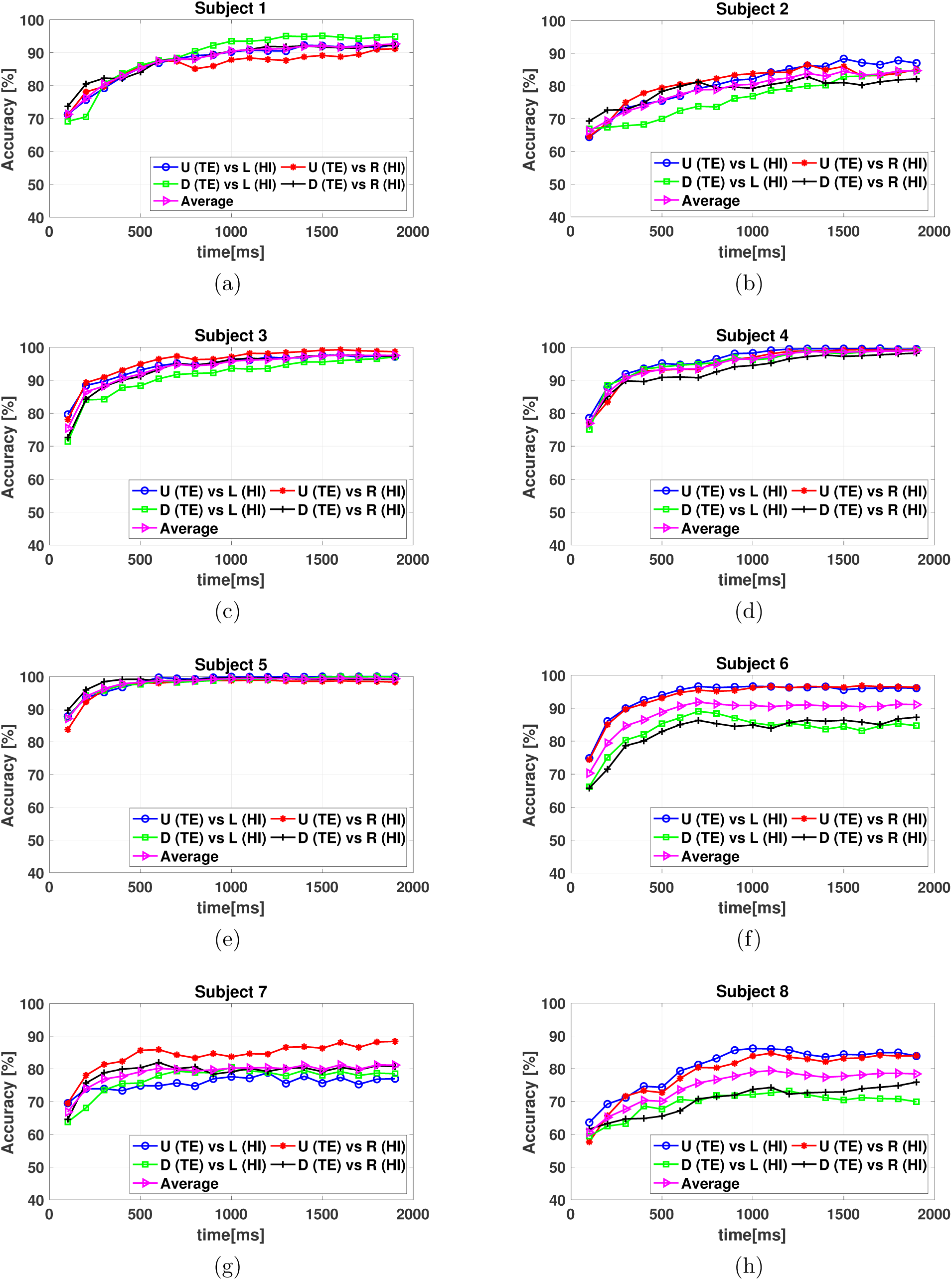
(a)-(h): Classification accuracy of tongue movement execution (TE) in up (U) or down (D) directions vs imagery (HI) of left (L) or right (R) hands for all subjects.

Considering these plots, it can be concluded that for most subjects, using the Table 5: Dataset 1-Classification accuracy results averaged across all subjects, for tongue movement execution (TE) vs tongue imagery (TI), and for tongue movement execution (TE) vs hand movement imagery (HI), at different post-stimulus instants. proposed approach, the accuracy reaches to its high levels within 500 − 1000 ms, except for Subject 5, for whom the highest accuracy was achieved earlier than 500 ms.

**Table 5:** Dataset 1-Classification accuracy results averaged across all subjects, for tongue movement execution (TE) vs tongue imagery (TI), and for tongue movement execution (TE) vs hand movement imagery (HI), at different post-stimulus instants.

| Duration (ms) | 300 | 500 | 700 | 1000 | 1500 | 1900 |
| --- | --- | --- | --- | --- | --- | --- |
| TE vs TI | $78.41 \pm 9.76$ | $82.27 \pm 9.23$ | $84.44 \pm 8.49$ | $85.39 \pm 8.66$ | $87.26 \pm 8.45$ | $87.35 \pm 8.39$ |
| TE vs HI | $82.14 \pm 9.37$ | $85.32 \pm 9.09$ | $87.64 \pm 8.64$ | $89.08 \pm 8.51$ | $90.01 \pm 8.32$ | $90.48 \pm 8.46$ |

To provide a comparison, the average accuracy results of the LSTM classifier across all subjects at different post-stimulus instants of 300, 500, 700, 1000, 1500, and 1900 ms are summarized in Table 5. These results suggest that using longer duration of recordings leads to some improvements in the classification accuracy.

### 4.3. Dataset 1: Significance of Considering the Dynamics of the Functional Connectivity Graphs on Classification Performance

To further emphasize the importance of including the dynamics contained within the sequence of the extracted features, we also considered a non-dynamic artificial neural network (ANN) classifier. In contrast to LSTM, the ANN classifier is incapable of taking advantage of the information conveyed by the sequencing of the feature vectors. We used an ANN model with two fully-connected hidden layers consisting of 20 neurons. The average classification results achieved at 500 ms using the ANN classifier for different classes of tongue movement execution vs imagery, and tongue movement execution vs hand movement imagery, compared to the accuracy results achieved from the LSTM classifier are presented in Figure 11. As can be seen, the average classification accuracy using the LSTM classifier is about 18% higher than the accuracy results achieved from the ANN classifier (across classes), which confirms that the inclusion of the information that lies in the temporal sequence of extracted features plays an important role in improving the performance.

**Figure 11:**
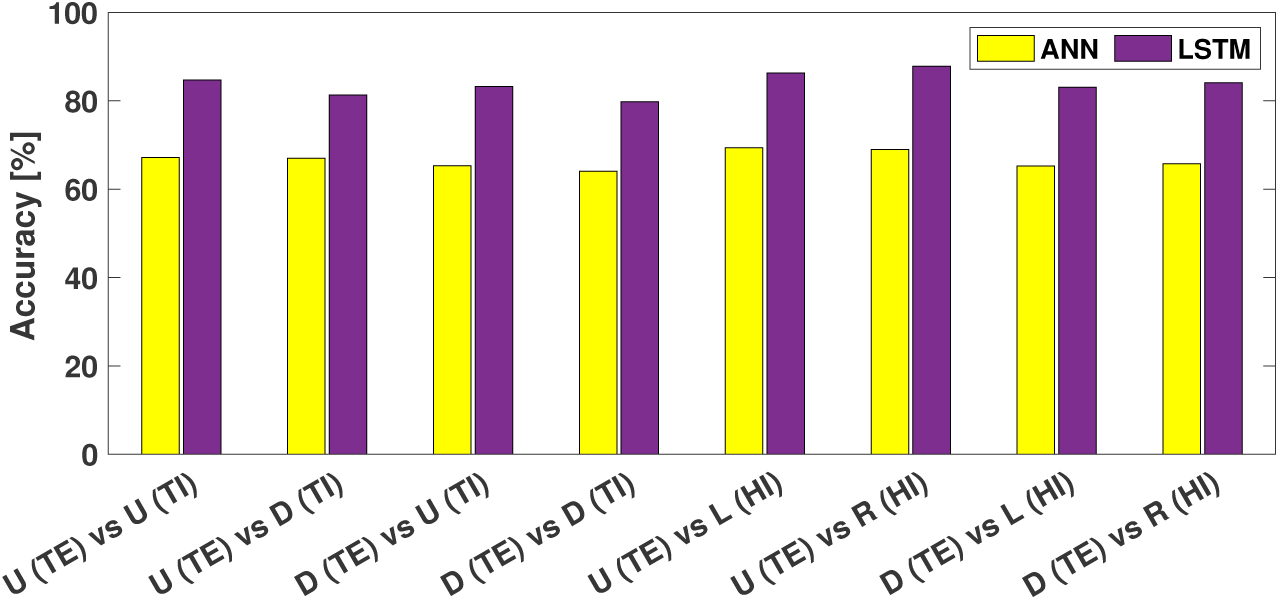
Average classification accuracy results, 500 ms after task onset, for different classes of tongue movement execution (TE) vs imagery (TI) and tongue movement execution (TE) vs hand movement imagery (HI) using ANN and LSTM classifiers.

Additionally, from Figure 12, one can see that, for the ANN-based classifier, the average classification accuracy does not improve as time goes by. This is caused by the exclusion of the temporal information of the sequence in the ANN classifier, thus, discriminating different tasks is only based on the spatial information of functional connectivity graphs. In summary, these results demonstrate the significance of considering the dynamics of the functional connectivity graphs for discriminating motor tasks.

**Figure 12:**
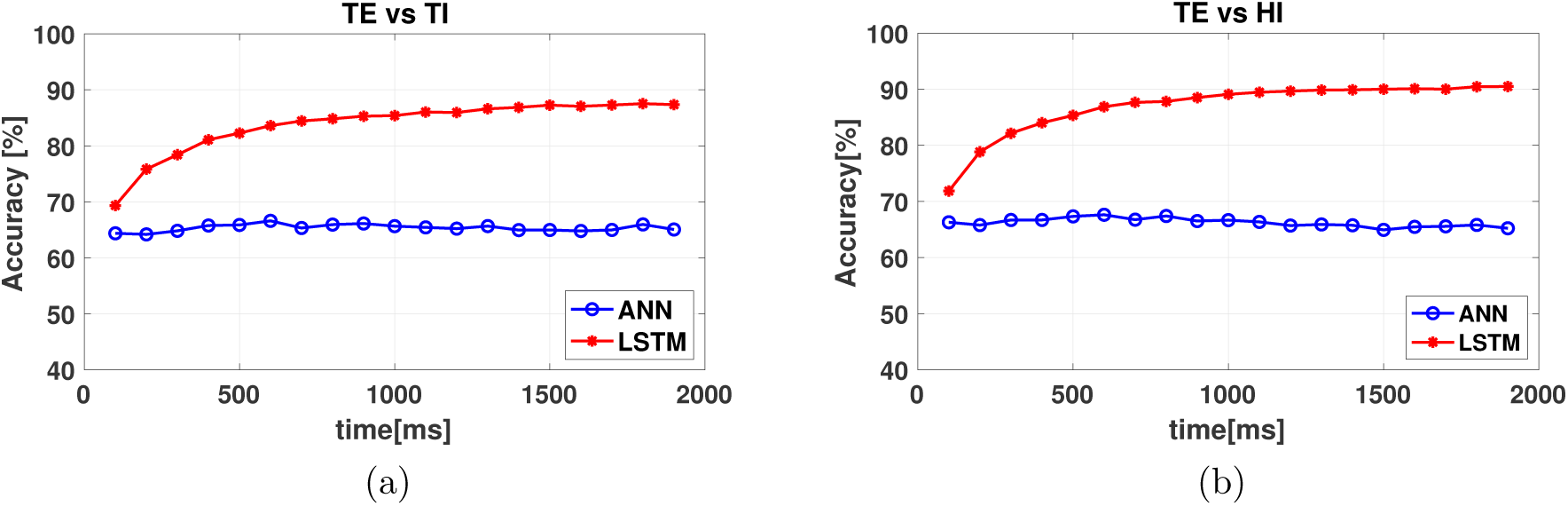
Average classification accuracy results for tongue movement execution (TE) vs (a): tongue movement imagery (TI) and (b): hand movement imagery (HI) using ANN and LSTM classifiers.

### 4.4. Dataset 2: Early classification of motor imagery tasks

In order to compare the performance of the proposed approach with existing methods, we also applied the classification algorithm to the BCI Competition IV dataset Iia (Dataset 2), which has been commonly used in BCI studies. Features extracted from the 3 s duration of motor imagery task ([3 − 6] s interval, see Figure 6(a)) were employed for training and validation. For testing, the time bin ending 800 ms, after the motor imagery task onset, was observed. The reason for selecting this duration is that when plotting accuracy graphs similar to Figures 9 and 10 for this dataset, we observed that around 800 ms a reasonable accuracy can be achieved. The classification results for differentiating various pairs of MI tasks (*i*.*e*. left hand vs right hand (L vs R), left hand vs both feet (L vs F), left hand vs tongue (L vs T), right hand vs both feet (R vs F), right hand vs tongue (R vs T), and both feet vs tongue (F vs T)) for all subjects are presented in Table 6. Accuracy results greater than 80% are also emphasized in bold.

**Table 6:** Classification accuracy results for classifying various pairs of left hand (L), right hand (R), both feet (F), and tongue (T) movement imagery tasks, 800 ms after the task onset for Dataset 2 (Dataset IIa from BCI competition IV).

| SUB | L vs R | L vs F | L vs T | R vs F | R vs T | F vs T |
| --- | --- | --- | --- | --- | --- | --- |
| A01 | 59.91 ± 6.88 | **80.62** ± 6.45 | **86.12** ± 4.92 | **81.49** ± 6.95 | **91.56** ± 4.46 | 55.18 ± 6.36 |
| A02 | 56.24 ± 6.62 | 72.87 ± 7.07 | 68.56 ± 7.45 | 75.35 ± 6.66 | 74.06 ± 6.92 | 60.39 ± 6.25 |
| A03 | 77.26 ± 6.01 | 76.08 ± 6.88 | **86.47** ± 5.24 | **86.21** ± 5.37 | **92.19** ± 3.70 | 70.57 ± 6.95 |
| A04 | 58.88 ± 7.48 | 63.23 ± 7.01 | 64.91 ± 6.98 | 68.84 ± 8.17 | 63.59 ± 6.53 | 60.92 ± 7.61 |
| A05 | 56.88 ± 6.39 | 52.03 ± 4.84 | 53.49 ± 5.96 | 53.44 ± 5.77 | 55.07 ± 6.36 | 51.03 ± 5.86 |
| A06 | 60.42 ± 8.21 | 60.67 ± 6.98 | 62.31 ± 7.24 | 56.23 ± 7.56 | 60.21 ± 7.90 | 58.59 ± 7.12 |
| A07 | 59.48 ± 6.06 | **88.13** ± 4.98 | **85.97** ± 5.54 | **83.22** ± 5.29 | **84.80** ± 6.43 | 72.74 ± 6.53 |
| A08 | **82.50** ± 5.44 | 69.69 ± 7.50 | **85.81** ± 5.48 | 76.91 ± 6.91 | **81.00** ± 5.85 | 76.90 ± 5.75 |
| A09 | **84.87** ± 5.64 | **86.04** ± 5.72 | **94.13** ± 3.69 | 65.82 ± 6.75 | 74.31 ± 7.09 | **80.66** ± 6.60 |
| Average | 66.27 ± 6.68 | 72.19 ± 6.45 | 76.42 ± 5.95 | 71.94 ± 6.67 | 75.20 ± 6.26 | 65.22 ± 6.59 |

As can be seen, the highest accuracy results have been achieved for left hand vs tongue, and for right hand vs tongue classification cases, whereas the lowest accuracy was obtained for feet vs tongue, and left hand vs right hand classification cases. One possible explanation for this observation is that the spatial and temporal patterns of the estimated functional networks are more discriminatory among some pairs of imagery tasks than others. Further exploration of the characteristics of these networks can deepen our understanding of the reasons for achieving different decoding accuracy results across different pairs of motor imagery tasks. Moreover, similar to the results obtained from Dataset 1, the results for this dataset were also subject-dependent. The accuracy results for subjects 1, 3, 7, 8, and 9 were higher than others, while the accuracy results for subjects 5 and 6 were mostly low. Although the accuracy results for other subjects were satisfactory, inclusion of these two subjects, has reduced the overall obtained averaged accuracy across subjects.

## 5. Discussion

There has been significant interest in studying task-based or resting-state brain’s functional connectivity in healthy and patient subjects, which has resulted in new understanding about the brain function as well as the mechanisms underlying brain-related disorders [117, 118, 119, 120, 121, 122, 123, 124, 125, 17, 126]. In BCIs, functional connectivity measures have been employed as features to discriminate various tasks [28, 26, 27, 22, 23]. This approach is in contrast to most of the BCI algorithms in which features are typically extracted from individual EEG electrodes. Previous studies suggest that using features from individual EEG electrodes may not provide enough information to discriminate motor tasks in specific groups of patients with motor disabilities [127]. For example, it has been shown that there exist differences in the band power patterns corresponding to motor imagery tasks across stroke patients with different location of lesion [128, 129, 130], and therefore, combining various features have been suggested as a solution to improve the classification performance [129]. Since performing tasks relies on the interactions among various brain regions, it is intuitive to consider functional connectivity-based characteristics rather than using features from isolated regions/electrodes for the purpose of discriminating intended tasks.

Brain is known to be a dynamical system in which interactions among different regions are time-varying [131, 17, 132, 133, 134]. These interactions are transient and rapid (*i*.*e*. established on the millisecond time scale). Accordingly, it has been suggested that studying the dynamics of functional connectivity networks provide a better understanding of the brain function compared to studying the brain in a static framework. Based on these facts, in this paper, we presented a novel feature extraction/classification framework which utilizes dynamic functional connectivity networks to decode the motor execution/imagery tasks. Furthermore, we aimed to reduce the time interval required to achieve reasonable accuracy.

The proposed framework includes segmentation of the EEG data into variable-length time intervals where the functional connectivity networks remain quasi-stationary. It is worth mentioning that most of the previously-presented methods (such as dynamic time warping (DTW), or microstates) for studying the brain dynamics through EEG recordings, have been based on changes in the activities that form on the scalp (sensor space)[17, 135, 136, 137]. In contrast, our novel source-informed segmentation algorithm identifies segment boundaries based on changes in the spatial characteristics of the functional networks in the cortex (i.e. the source space). This makes the algorithm informed by the source space, without requiring computationally-complex source localization algorithms [101].

In the proposed framework, the sequence of extracted connectivity features are passed to the LSTM classifier to take advantage of the information that lies in the temporal sequence of the constructed connectivity graphs. The obtained classification accuracy results for Dataset 1 indicate that the proposed method can successfully discriminate tongue movement execution vs tongue (82.27%)/hand (85.32%) movement imagery tasks at only 500 ms after the task onset. These results suggest that the connectivity patterns of motor execution/imagery tasks along with their temporal dynamics can provide sufficient information for solving the classification problem within a short interval of in the order of hundreds of milliseconds. Therefore, the proposed feature extraction/classification framework can be utilized to reduce the required buffering lag for the BCIs, and thereby, increasing their speed. It should be noted that the time instants indicated for the two datasets (i.e. 500 ms for Dataset 1, and 800 ms for Dataset 2) were selected as examples to show that by using the proposed method, good classification accuracy can be obtained as early as these time instants after the task onset. However, as illustrated in Figures 9 and 10 due to the dynamic nature of the proposed approach, the decision can be generated at any time instant since the first identified segment. This is the feature offered by the proposed method that we refer to as “early decoding”.

It is worth to mention that for Dataset 1, we have considered tongue movement execution as one of the tasks for the BCI. The importance of discriminating tongue movement execution vs other imagery tasks is that the movement of tongue is often available even in patients with severe motor disabilities. Furthermore, tongue is a relatively strong muscle and can be moved in various directions [138]. If a patient possesses the ability of moving his/her tongue, the BCI can actually benefit from this available physiological signal. Therefore, in cases where tongue movement is available from the patient, it can be used along with other motor imagery tasks to increase the number of control commands.

Our results suggest that the temporal dynamics of functional networks play key roles in early decoding capability offered by the proposed framework. This was highlighted by comparing the classification results achieved from two classifiers: the LSTM which considers both the temporal dynamics and the spatial information of functional connectivity graphs, and ANN which only utilizes the spatial information. It was observed that for the ANN classifier, there was not much difference in the obtained classification accuracy results along time, while in the case of the LSTM classifier, the average accuracy results improved along the time spent after task onset (Figure 12). These results further confirm the importance of the inclusion of the temporal information of the functional connectivity networks in achieving good classification performance within a short interval. The significance of including the temporal information can be observed as early as 100 ms after the task onset, where the average classification accuracy using LSTM is ∼ 5% higher than the case of using ANN classifier. This difference increases to 16.39 − 17.99% and 22.28 − 25.26% for 500 and 1900 ms after the task onset, respectively. It can be concluded that the interactions among the brain regions and how they evolve over time contribute to discriminatory information needed to solve the classification problem.

The proposed method was employed to differentiate motor execution vs imagery tasks. The similarities and differences in connectivity patterns of motor execution and imagery tasks were investigated in [139, 140, 141, 123].

For example, in [139], coupling patterns among occipital and motor regions in the beta frequency band were reported to be different during hand movement execution and imagery. In [140], it was shown that the key nodes in netowrks related to motor execution and imagery tasks are located in different areas. In [141] and [123] functional connectivity analysis based on ERS/ERD and phase synchronization showed similar connectivity patterns among contralateral brain regions for movement execution and imagery of finger tapping [141], and foot and hand [123]. In [142], the results of effective connectivity networks associated with finger tapping execution and imagery indicated that the coupling strength of the feedforward network from dorsolateral prefrontal cortex to premotor cortex was greater during motor execution tasks than to motor imagery tasks, whereas the coupling strength of the feedforward network from premotor cortex to supplementary motor area and the feedback network from the primary motor cortex to premotor cortex were higher for motor imagery tasks. In [143], it was suggested that increases in *β*-band connectivity occurs similarly in both movement execution and imagery tasks, while in *µ* band, motor execution and imagery tasks are associated with different connectivity patterns. Considering the results of these studies, it can be concluded that depending on the analysis methods and variables that are selected as the basis of comparisons, one might observe common or different connectivity patterns among motor execution and imagery tasks. In this paper, we showed that using the proposed method, *dynamic* functional connectivity networks can provide discriminatory information corresponding to motor execution vs imagery tasks. Frequency-specific analysis can bring more insights into understanding how different EEG frequency bands contribute in differentiating motor execution vs imagery tasks. We aim to investigate this problem in our future studies.

Table 7 summarizes the classification results for some of the recent work that have used Dataset 2 along with the required duration of the data that was used to achieve these classification accuracy results. It can be seen that, our work shows the shortest buffering lag (800 ms) while other methods required buffering lags equal or greater than 2 s to achieve the reported accuracy results.

**Table 7:** Comparison of classification accuracy results and the required time, of the proposed method and existing work using Dataset 2 (Dataset IIa from BCI competition IV)

| Ref. | Year | Duration (s) | Feature Type | Classifier | Accuracy (%) |  |  |  |  |  |
| --- | --- | --- | --- | --- | --- | --- | --- | --- | --- | --- |
|  |  |  |  |  | L vs R | L vs F | L vs T | R vs F | R vs T | F vs T |
| [37] | 2016 | 3 | Channel-based | SVM | 74.92 | - | - | - | - | - |
| [25] | 2016 | 3.5 | Functional Connectivity | SVM | 64.27 | - | - | - | - | - |
| [80, 38] | 2017 | 3 | Channel-based | RBM | 64.60 | 65.60 | 61.90 | 66.80 | 70.20 | 64.60 |
| [144] | 2017 | unknown | Channel-based | MNN | 79.52 | 83.19 | 84.64 | 81.44 | 83.40 | 78.64 |
| [52] | 2018 | 2 | Channel-based | SVM | 82.50 | - | - | - | - | 84.00 |
| [42] | 2018 | 2 | Channel-based | MDRM | 79.93 | 85.50 | 84.30 | 85.43 | 83.75 | 74.78 |
| [38] | 2018 | 3 | Channel-based | LSTM | 74.80 | 74.20 | 72.50 | 76.30 | 74.30 | 79.60 |
| [55] | 2019 | 2 | Channel-based | MDRM | 80.98 | - | - | - | - | - |
| [23] | 2019 | 3 | Functional Connectivity | LS | 71.00 | 74.00 | 74.00 | 71.00 | 74.00 | 65.00 |
| This work | - | **0.8** | Functional Connectivity | LSTM | 66.27 | 72.19 | 76.42 | 71.94 | 75.20 | 65.22 |

Due to differences in the preprocessing steps, types of extracted features, computational complexity, and choice of classifiers, a fair comparison is not possible across different work. Another main difference that should be pointed out is that except ours, the methods in Table 7 cannot be used for early decoding purposes. That is, they would require the processing of the full considered interval of EEG data, before coming up with a decision, while our method offers the possibility of decoding the tasks as early as the first identified segment, due to it dynamic (segment-by-segment) classification nature. In terms of types of extracted features, [23] and [25] are the closest to our work for comparison as they have employed functional connectivity-based features, albeit without considering the dynamics. Compared to these two studies, it can be observed that our work has led to better accuracy results for majority of listed MI tasks, while requiring only 800 ms after task onset, further suggesting the importance of the inclusion of dynamic information for discriminating motor imagery tasks. In terms of classifiers, [38] has also used LSTM, however, the extracted features are time-domain and channel-based. The reported classification accuracy results from this method for duration of 3 s after task onset are comparable to our results after 800 ms. Furthermore, the results for [38] which uses a deep learning classification algorithm based on Restricted Boltzmann Machines (RBM) [80], are lower for all listed pairs of MI tasks, as compared to our results. In [144], a method for reducing the effects of noisy trials was used, which has resulted in improved classification performance. In our case, we had only considered basic preprocessing steps (filtering and ICA). The possibility of achieving higher accuracy results with more advanced preprocessing methods such as those proposed in [144] requires further investigation. Also, the buffering lag for using this result was not directly reported in [144], however, it seems the duration of the trial was used in the analysis, which is considerably longer than what we used in our work. The works [37, 52, 42, 55] have used channel-based features extracted from 2 or 3 s duration. Despite the good classification results reported in [37], implementing this method is not computationally efficient due to the procedures involved in the decoding algorithm. In [52], both the filter bandwidth and the time window used for feature extraction were optimized for accuracy. In [42] features are extracted from specific EEG frequency bands. In our work, we did not consider band-specific results, and it would be interesting to investigate whether optimizing bandwidth or employing frequency-specific dynamic functional connectivity networks as features will improve the accuracy results. Finally, in [55], a Riemannian-based approach is used which incorporates inter-subject data to reduce the dimensionality of the covariance matrices through regularization techniques. Including the inter-subject data in our proposed dynamic functional connectivity-based method can be an interesting topic for the future studies. In summary, it can be concluded that although some of the studies in Table 7 have reported higher classification accuracy results as compared to our case, we have shown that within a notably shorter duration, reasonable classification accuracy can be achieved using our proposed method. Inclusion of advanced-preprocessing techniques, or optimizing the performance based on band selection, could further improve our accuracy results. Our early decoding method is well-suited for applications that require decoding the user’s intentions within a short interval after the onset of the task, while other methods could be more preferable if higher decoding accuracy is required and the duration of the buffering lag is not a concern.

To the best of our knowledge, this is the first study that demonstrates the possibility of early decoding of motor tasks in BCIs from EEG recordings. There are a few limitations in this study. Here, we have performed off-line processing and classification of EEG data. In order to use this method in real-time BCIs, a framework for a real-time implementation of the feature extraction algorithm should be developed. With the current progress in the development of computational hardware and processing units, this seems to be achievable in the near future. Moreover, previous studies on MI-based EEG classification [145, 146, 147, 58] have demonstrated the role of specific EEG frequency bands in the classification of MI tasks. We did not incorporate frequency decomposition here. An extension of this work could be focused on the extraction of functional networks from different EEG frequency bands to investigate the significance of each frequency band in addressing the early decoding problem. Additionally, we only considered basic FIR filtering and artifact removal using ICA. The performance of the proposed method can further be improved by incorporating noise and artifact reduction techniques in the algorithm and/or utilizing mechanisms to detect and remove noisy trials as suggested in [144].

It is also worth mentioning that although we have verified the early-decoding effectiveness of the proposed approach using two different datasets, the collected data belonged to healthy groups. For the purpose of implementing BCI-based assistive technologies for patients, testing the proposed framework on data collected from patients would be necessary. Moreover, in this work, we considered the decoding problem for movement-related tasks. However, the proposed framework can also be applied to other tasks, such as P300 and cognitive tasks, to evoke brain activities in BCIs. Implementing the proposed method for early decoding of other mental activities from EEG recordings may offer the ability to improve the speed and practicality of the BCIs that are based on these tasks.

## 6. Conclusions

This paper introduced a new feature extraction method based on dynamic functional connectivity networks for early decoding of EEG signals. The proposed method comprised of two steps: first, segmenting the EEG data into quasi-stationary temporal blocks during which functional networks sustain their connectivity, and second constructing functional connectivity graphs for each identified segment. An LSTM classifier was then employed for the classification due to its advantageous utilization of the contained memory cell which allowed for processing a sequence of features.

The proposed method enabled us to differentiate among tongue movement execution vs tongue or hand movement imagery tasks within a short interval in the order of hundreds of milliseconds (e.g. 500 ms) with good accuracy. These results indicated, for the first time, that the required duration of EEG data on a given trial for decoding motor execution and imagery tasks, can be significantly reduced compared to existing methods, suggesting that this framework is an efficient approach for improving the speed of BCIs. Additionally, we showed that the early decoding capability relied on both spatial and temporal information of functional connectivity networks that were captured by using the proposed feature extraction method and utilizing the LSTM classifier.

This study was the first step in addressing the early classification problem from EEG data. The extension of the proposed method for band-specific early decoding, and multi-class problems are considered as our future work.

## Acknowledgment

The authors would like to acknowledge the support from NSF (Award 1841087), Siemens, and DARPA. The authors would also like to thank Jennifer Huang for her help in EEG data collection.

## Appendix

**Table A1:** List of Acronyms.

|  |  |
| --- | --- |
| AFNN | Adaptive Fuzzy Neural Network |
| ANN | Artificial Neural Network |
| BCI | Brain Computer Interfaces |
| BPNN | Back-Propagation Neural Network |
| BTSPRT | Balanced Threshold Sequential Probability Ratio Test |
| CPMS-CNN | Channel-Projection Mixed-Scale Convolutional Neural Network |
| CNN | Convolutional Neural Network |
| CSP | Common Spatial Pattern |
| DNN | Deep Neural Network |
| DPL | Dictionary Pair Learning |
| EEG | Electroencephalography |
| ELM | Extreme Learning Machine |
| EOG | Electrooculography |
| ERD | Event-Related De-synchronization |
| ERS | Event-Related Synchronization |
| FIR | Finite Impulse Response |
| FLDA | Fisher Linear Discriminant Analysis |
| GA | Genetic Algorithm |
| GB | Gradient Boosting |
| GSLDA | Generalized Sparse Linear Discriminant Analysis |
| HMM | Hidden Markov Model |
| HS-CNN | Convolutional Neural Network with hybrid Convolution Scale |
| ICA | Independent Component Analysis |
| IT2FLS | Interval Type-2 Fuzzy Logic System |
| kNN | k-Nearest Neighbors |
| KSVM | Kernel Support Vector Machine |
| LDA | Linear Discriminant Analysis |
| LR | Logistic Regression |
| LRP | Layerwise Relevance Propagation |
| LS | Least-Square |
| LSTM | Long Short Term Memory |
| MDRM | Minimum Distance to Riemannian Mean |
| ME | Motor Execution |
| MI | Motor Imagery |
| MNN | Modified Neural Network |
| NB | Naive Bayse |
| NBPW | Naive Bayesian Parzen Window |
| NN | Neural Network |
| pCNN | ; Spectrogram-based Convolutional Neural Network |
| PCSP | Probabilistic Common Spatial Pattern |
| PSO | Particle swarm optimization |
| RANFN | Recurrent Adaptive Neuro-Fuzzy Network |
| RBM | Restricted Boltzmann Machines |
| RCNN | Recurrent Convolutional Neural Network |
| RF | Random Forest |
| SBL | Sparse Bayesian Learning |
| SRC | Sparse Representation-based Classification |
| SRDA | Spectral Regression Discriminant Analysis |
| RVM | Relevance Vector Machine |
| SVD | Singular Value Decomposition |
| SVM | Support Vector Machine |
| SUSS-SRKDA | Sequential Updating Semi-Supervised Spectral Regression Kernel Discriminant Analysis |
| WNB | Weighted Naive Bayesian |

## Footnotes

‡ Note that due to high volume of papers in this field, we only considered highly-cited studies that considered classification of motor-related tasks, and were published after 2014.

